# Planar cell polarity aligns epithelial migration to coordinate tissue architecture and functional zonation

**DOI:** 10.1101/2025.10.30.685585

**Authors:** Rebecca F Lee, Dami Oluwatade, Kaelyn Sumigray

## Abstract

Epithelial tissues must coordinate collective migration with functional specialization to maintain organ integrity, but the mechanisms linking these processes remain unknown. Here, we define the planar cell polarity (PCP) pathway as a master coordinator that translates molecular asymmetry into tissue-scale organization during postnatal intestinal development. We show that PCP drives the developmental transition from random to linear epithelial migration by aligning actin-based basal protrusions and polarizing basement membrane integrins. This creates coherent cellular streams that form characteristic villus ribbons. Strikingly, we discover that disrupting this migration coherence directly perturbs functional zonation, as loss of PCP causes disordered migration that expands specialized metabolic domains beyond their spatial boundaries. These findings reveal a previously unrecognized principle of epithelial organization: directional collective migration actively maintains functional compartmentalization, establishing migration pattern as a fundamental determinant of how tissues preserve specialized function during continuous turnover.

## Introduction

During development, morphogenic events build upon one another to sculpt the organization of organs so that their final organization reflects their physiological function. The mammalian small intestine is a striking example of this principle: distinct yet spatially integrated morphogenic programs converge to generate the stereotyped crypt-villus architecture that maximizes absorptive surface area and establishes distinct biochemical microenvironments along the epithelial axis. The mature epithelium consists of proliferative crypts, which house stem and progenitor cells, and absorptive villi, which are populated by differentiated enterocytes and secretory cells. Each villus is fed by approximately 10-12 crypts^1^, whose progeny migrate along defined linear trajectories to form sharply bounded clonal ribbons of cells that extend from the crypt-villus junction to the villus tip^2,3^. Although this pattern is well described in adult tissue, the developmental mechanisms that establish and maintain villus ribbons remain largely unknown. This organization is not incidental: the linear architecture of villus ribbons preserves spatial segregation of specialized gene expression domains^4^, ensuring that distinct regions of the villus perform unique functions in nutrient processing^5,6^.

While embryonic organogenesis has been studied in depth, the intestine continues to undergo rapid and extensive remodeling after birth. During this postnatal period, the crypt-villus axis is established^7^, crypts become monoclonal^2^, and villus architecture matures to its final pattern^1^. Daughter cells from each crypt then migrate upward in continuous stripes, producing the characteristic villus ribbons of the adult epithelium. Previous studies have attributed ribbon formation to the stochastic fixation of crypt clonality^2,3^, without considering an active mechanism for maintaining linear trajectories. However, the sharp boundaries between neighboring ribbons and the persistence of these boundaries during epithelial turnover imply active coordination of migration direction. How villus cells collectively align their movement, and why such linear migration paths are maintained, remains unknown.

The intestinal epithelium renews more rapidly than any other mammalian tissue. Cells migrate from the base of the crypt to the tip of the villus in 3-7 days before being extruded.

Historically, this flow was modeled as a passive “conveyor-belt” in which mitotic pressure from crypt proliferation displaces differentiated cells upward^8–10^. More recent work has challenged this model, revealing that epithelial cells actively migrate through the formation of polarized actin-based basal protrusions that generate traction along the basement membrane^11^. These findings raise a fundamental question: when and how do villus cells acquire directional polarity and organize into linear, non-overlapping ribbons?

Here, we define the developmental emergence of directional migration and reveal how it is coordinated along the intestinal epithelium. We find that during postnatal villus maturation, migration paths become progressively linear as actin-based basal protrusions and basement membrane integrins become increasingly polarized. This cellular-level organization depends on the planar cell polarity (PCP) pathway, which we identify as a critical regulator of epithelial migration directionality and mechanical coherence in the postnatal intestine. Loss of PCP disrupts protrusion alignment and integrin localization, producing disordered migration trajectories, loss of linear villus ribbons, and local architectural defects. Moreover, the disruption of migration coherence leads to expansion of villus tip gene and metabolic domains, revealing that PCP-mediated alignment of epithelial migration is necessary to preserve spatial zonation of function.

Altogether, these findings identify PCP as a developmental organizer that links planar polarity, integrin-mediated adhesion, and collective migration to maintain the spatial fidelity of intestinal architecture and function. Through this integration, local molecular asymmetries are translated into directional epithelial motion, ensuring that cellular turnover, morphogenesis and nutrient-processing zonation remain coupled along the crypt-villus axis.

## Results

### Directional epithelial migration emerges during postnatal villus morphogenesis

To define when epithelial cells begin to migrate in linear paths, we first visualized clonal cell populations along the villus axis and assessed their ability to form linear ribbons throughout postnatal development. Tamoxifen-induced lineage tracing from Lgr5-EGFP-IRES-Cre^ER^; mTmG mice at postnatal day (P)0-P1 produced sparse GFP-labeled clones in stem cells and their progeny (Fig 1A). At P10, clones had irregular, jagged boundaries that crossed neighboring territories, suggesting uncoordinated migration. Over the course of 10 days, clones began to adopt more linear shapes, and by P20, they had resolved into sharply bounded ribbons extending from the villus base to the tip (Fig 1B). Quantification of boundary “linearity”, the ratio of the traced boundary length to Euclidean distance, confirmed a progressive decrease from ∼1.35 at P10 to 1.05 at P20 (Fig 1C), approaching the geometry of a straight line. This transition marks the establishment of reproducible, directionally coherent migration trajectories coincident with villus maturation.

**Figure 1.**
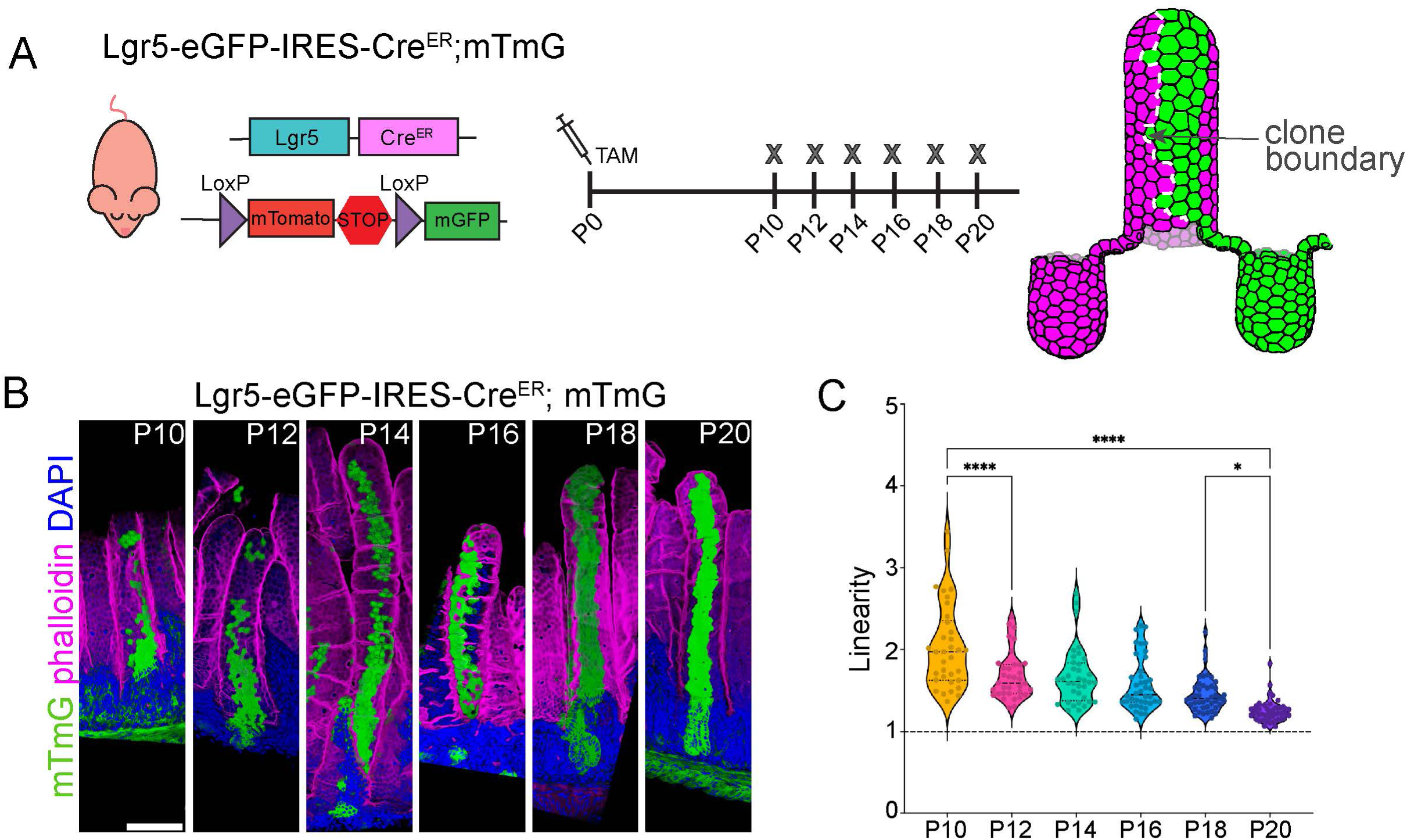
Directional epithelial migration emerges during postnatal villus morphogenesis. A) Schematic of mouse model and tamoxifen administration scheme and measurement of clonal boundary. Dashed line represents clone boundary. B) Representative villi from P10 through P20 intestines showing GFP-labeled clones extending along the villus axis. Scale bar, 100 µm. C) Quantification of ribbon linearity, measured as the ratio of traced boundary length to Euclidean distance. *\* p* < 0.05, **** *p* < 0.0001.

### Collective migration becomes coherent and processive with villus maturation

To directly visualize this dynamic transition directly, we performed long-term live imaging of 2D enteroid monolayers derived from P10 and P20 intestines (Fig 2A-F). P10 cells exhibited displayed short, erratic trajectories (Fig 2A-C) with minimal net displacement (Fig 2G). In contrast, cells within P20 monolayers displayed broad, coordinated streams of unidirectionally migrating cells (Fig 2D-F). Notably, P10 cells exhibited minimal productive movement (Fig 2J), as shown by the representative tracks (Fig 2B) and displacement over 14 h (Fig 2G). In contrast, cells within P20 monolayers efficiently moved (Fig 2D-J) in linear paths (Fig 2G-I).

**Figure 2.**
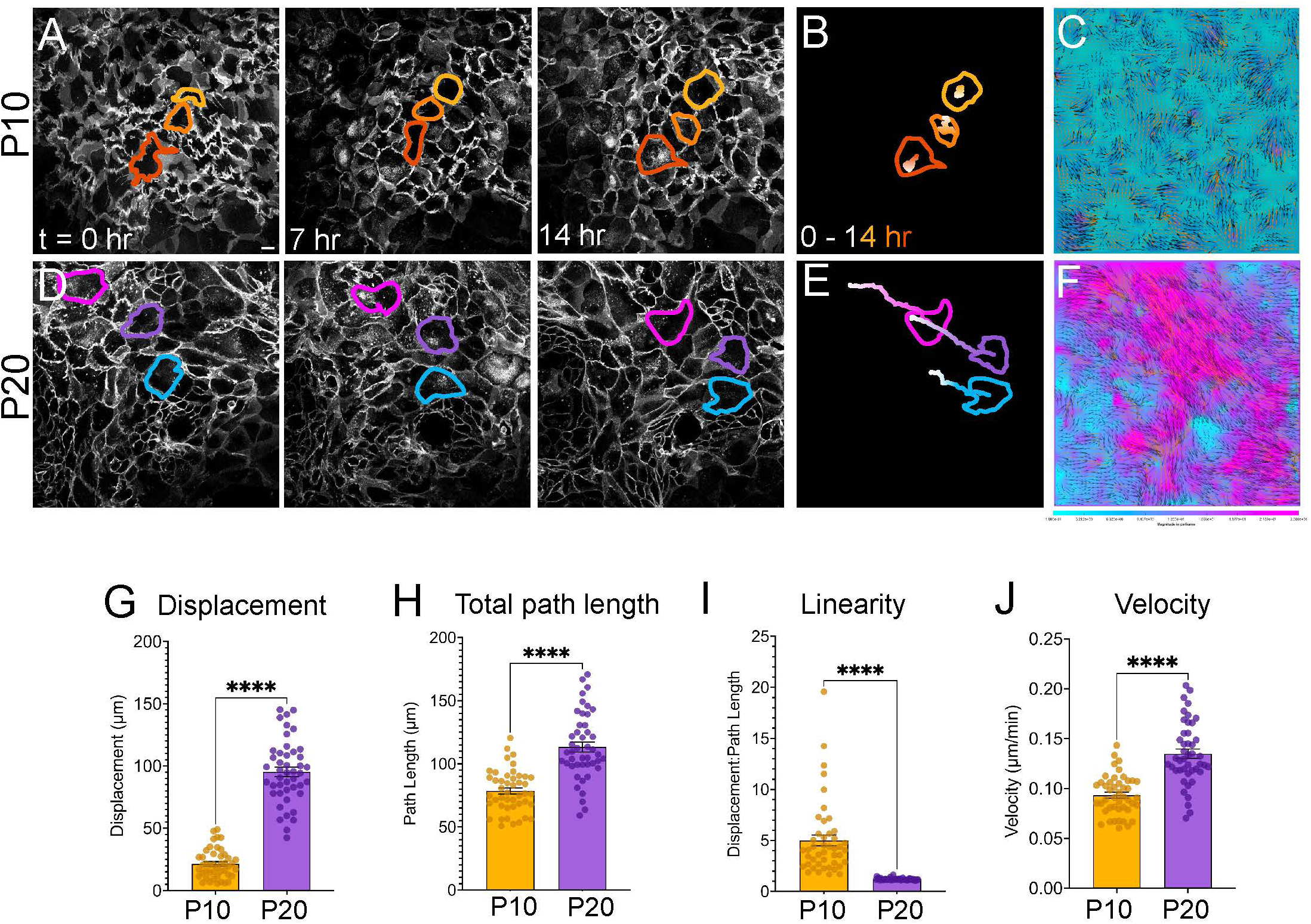
Collective epithelial migration becomes coherent and processive with villus maturation. A) Time series of 2D enteroid monolayer live imaging from P10 mTmG intestine. B) Schematic of cell paths from A. C) Particle image velocimetry (PIV) of P10 monolayer showing slow uncoordinated movement. D) Time series of 2D enteroid monolayer live imaging from P20 mTmG intestine. E) Schematic of cell paths from D. F) PIV of P20 monolayer showing rapid collective flow. G-J) Quantification of displacement, total path length, linearity and velocity of tracked cells from P10 and P20 enteroid monolayers. At least 45 cells were analyzed per sample, N = 3 mice per genotype. ****, *p* < 0.0001.

Particle image velocimetry and linearity measurements revealed that collective motion evolved from random to collective streams (Fig 2C, F). These data establish that postnatal villus maturation is accompanied by a fundamental shift in cell behavior, from stochastic individual movements to a coordinated, tissue-scale migration program.

### The planar cell polarity pathway is activated during the onset of directional migration

Having established when coherent migration trajectories arise, we next sought to identify the molecular machinery responsible for this developmental transition. Transcriptomic profiling of publicly available bulk RNA-sequencing data of postnatal intestinal epithelial cells^7^ from P6 to P10 identified strong induction of planar cell polarity (PCP) components, including the genes encoding the core transmembrane proteins Vangl1, Celsr1, Celsr2 and Celsr3 (Fig 3A-B). To study the expression and localization of PCP proteins in the postnatal intestinal epithelium, we utilized a transgenic Rosa26-Vangl2-GFP mouse line^12^. Vangl2-GFP was more highly expressed in crypts compared to villi at all time points examined (Fig 3C-F, K). Vangl2-GFP localization was initially diffuse at P5, becoming more enriched along crypt cell membranes by P15-P20 was diffusely localized throughout crypt cells (Fig 3C-K). The specific enrichment of Vangl2-GFP in crypts was unexpected, given that the transgene is driven by the ubiquitously active *Rosa26* promoter rather than a crypt-specific element. Consistent with this, most PCP transcripts showed comparable expression between crypts and villi at each time point (Fig 3A). Notable exceptions were *Vangl2* (1.92-fold higher expression in P10 crypts compared to P10 villi; *p* < 0.0001), *Celsr1* (1.55-fold higher; *p* = 0.007), and *Celsr2* (2.04-fold higher; *p <* 0.0001). Because loss of any single core PCP component destabilizes the remaining complex and prevents membrane localization^13^, these modest but coordinated enrichments may allow stable complex formation only within crypt cells. We propose that this localized assembly of PCP complexes generates a polarity field that orients epithelial migration as cells exit the crypt and ascend the villus.

**Figure 3.**
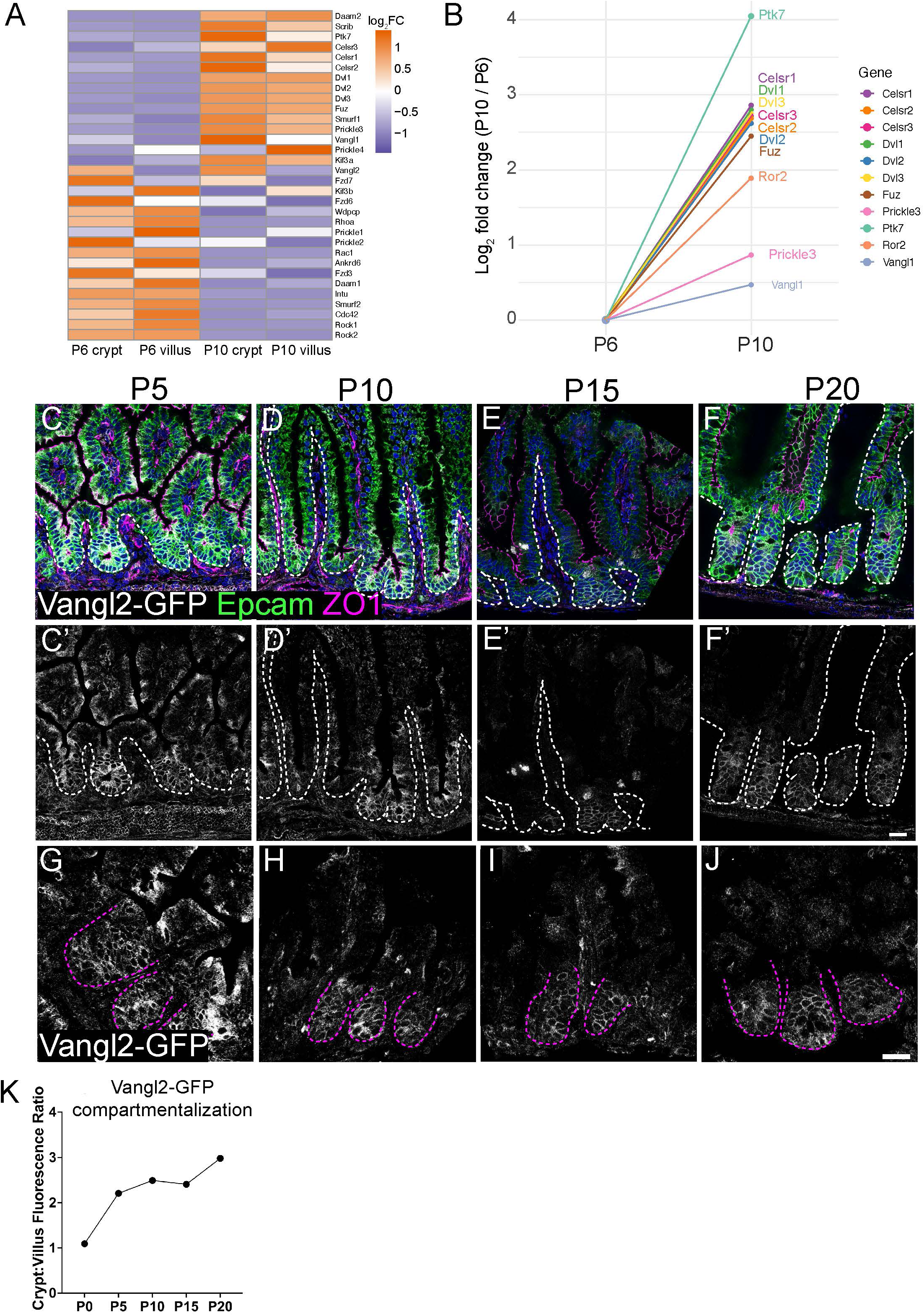
PCP signaling is induced during villus maturation and establishes a polarity field in crypts. A) Heatmap of PCP gene expression in postnatal intestinal epithelial cells from P6 and P10. B) Relative expression change (log2FC) from P6 to P10 of select PCP genes. C-F; C’-F’) Representative images of Rosa26-Vangl2-GFP reporter intestines at P5, P10, P15, and P20. Vangl2-GFP (white) becomes progressively enriched along crypt cell membranes. Epcam (green) marks epithelial cells, ZO1 (magenta) marks tight junctions. Dashed white line marks basement membrane. G-J) High magnification views of crypts showing Vangl2 enrichment on crypt cell membranes. Magenta dotted line outlines crypts. K) Quantification of relative Vangl2- GFP intensity between crypts and villi at each stage.

### Loss of PCP signaling disrupts migration dynamics

To determine whether the PCP pathway was involved in developmental acquisition of linear ribbon patterns, we induced the loss of either Vangl1/2 or Dvl2 in the intestinal epithelium using Villin-Cre^ER^. Recombination was induced at P2 to capture the developmental period preceding migration alignment. While we utilize two distinct cKO models, we have found that they completely phenocopy each other. To assess villus ribbon patterns, we utilized the multicolor iChr-mosaic reporter to clonally label nuclei^14^. By P20, control villi displayed sharply bounded, parallel ribbons (Fig 4A), whereas PCP cKO villi exhibited pronounced disorganization among clones, with tortuous ribbons that frequently intersected neighboring clones (Fig 4B). Quantitative analysis of ribbon identity across the villus surface confirmed that PCP loss abolishes the alternating patterning expected from a sharp linear boundary (Fig 4C-D). In controls, neighboring ribbons displayed near-perfect anti-correlation (frequency = +1.0 ± 0.1 vs. -1.0 ± 0.1) with a sharp transition between zones 4 and 5 – the location of the clonal boundary. In PCP cKO tissue, this periodic organization collapsed to near random levels: the mean frequency decreased to 0.4 ± 0.3. The amplitude and reproducibility of the ribbon frequency pattern dropped sharply in PCP cKO tissue, indicating a loss of coherent spatial organization. Despite this geometric disarray, crypt clonality rates and bulk turnover rates were unchanged, indicating that PCP loss specifically affects migration directionality rather than proliferation or homeostasis.

**Figure 4.**
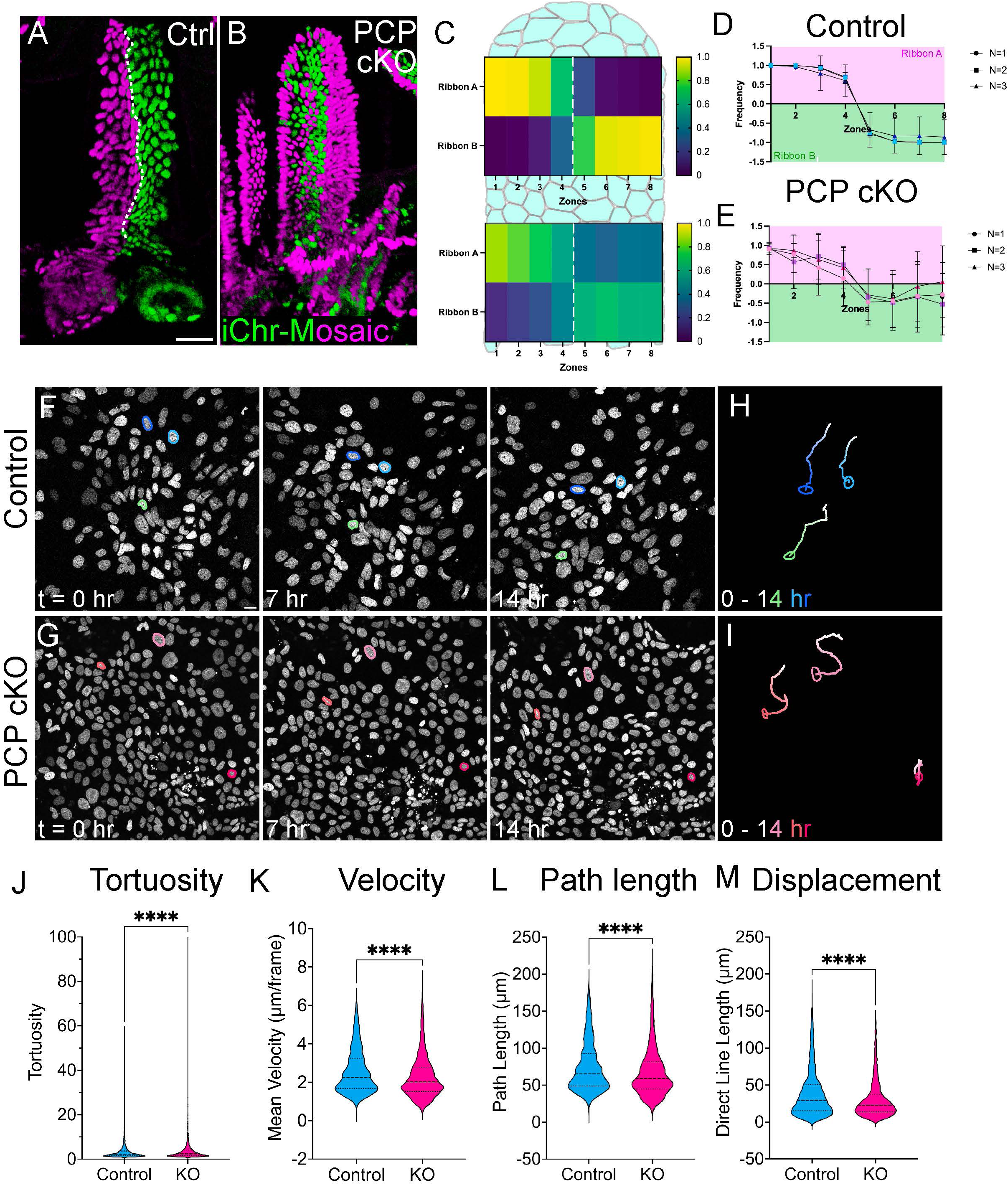
Loss of PCP disrupts epithelial migration dynamics and villus ribbon organization. A-B) Representative villi from control and Villin-Cre^ER^; Dvl2 fl/fl; iChr intestines analyzed at P20. In A) linear ribbon boundary is marked by dotted white line. C-D) Quantification of ribbon periodicity across the villus surface. In controls, neighboring ribbons exhibit strong anti-correlation, reflecting strong boundaries. PCP cKO villi display near-random organization, indicating a breakdown of spatial coherence. F-G) Time lapse imaging of 2D enteroid monolayers from control (F) and PCP cKO (G) intestines at P20. Scale bar, 20 µm. H-I) Schematic of cell tracks from F and G. J-M) Quantitative analysis of live cell trajectories. ****, *p* < 0.0001.

To definitively test whether PCP is required for cell migration dynamics, we performed time-lapse imaging of enteroid monolayers derived from P20 WT or PCP cKO intestines (Fig 4F- G). In controls, cell trajectories were long and highly aligned with a common axis, generating continuous flows (Fig 4H). In PCP cKO, trajectories were shorter, erratic and randomly oriented (Fig 4I). Analysis of live cell dynamics revealed that PCP cKO cell tracks were more tortuous (Fig 4K), less processive (Fig 4L), and slower (Fig 4M).

Together, these data establish that the PCP pathway is indispensable for converting stochastic postnatal migration into organized, linear movement. Without PCP, enterocytes still migrate, but do so incoherently, leading to distorted ribbon geometry and villus topology despite otherwise normal differentiation and renewal.

### PCP orients basal protrusions to organize linear villus ribbons

The loss of directional migration in PCP cKO villi suggested that disrupted ribbon geometry may arise from a failure of individual cells to coordinate their movement. Because epithelial migration in the intestine is mediated by actin-rich basal protrusions that orient in the direction of migration and generate traction along the basement membrane^11^, we hypothesized that protrusion orientation correlates with the emergence of linear ribbon patterns. To test this, we first examined protrusive behavior across postnatal development using high-resolution confocal imaging of individual cells within villi from P10 (before ribbon formation) and P20 (after ribbon formation) Lgr5- EGFP-IRES-Cre^ER^; mTmG intestines (as in Fig 1A). At P10, when ribbon boundaries were still irregular, enterocytes displayed an array of short, randomly oriented basal protrusions in all directions (Fig 5A-C). By P20, when villus ribbons appeared straight and continuous, nearly all basal protrusions were uniformly oriented along the migration axis (Fig 5D-F). Quantification across 3D reconstructions showed a clear developmental shift. The distribution of protrusion angles narrowed (Fig 5C, F), and the mean front-to-back surface area increased from ∼1 at P10, reflecting an even distribution of protrusions over the basal surface of the cell, to ∼3 by P20 (Fig 5G), reflective of the significant polarization of protrusions to the leading side of the cell. Interestingly, as protrusions became more polarized with age, the number of protrusions per cell decreased (Fig 5H). Therefore, polarization of basal protrusions correlated with the developmental timing of linear paths along villi. These data indicate that basal protrusion polarity emerges gradually during the developmental period when epithelial migration becomes coherent, suggesting that protrusive organization is a key structural correlate of villus ribbon maturation.

**Figure 5.**
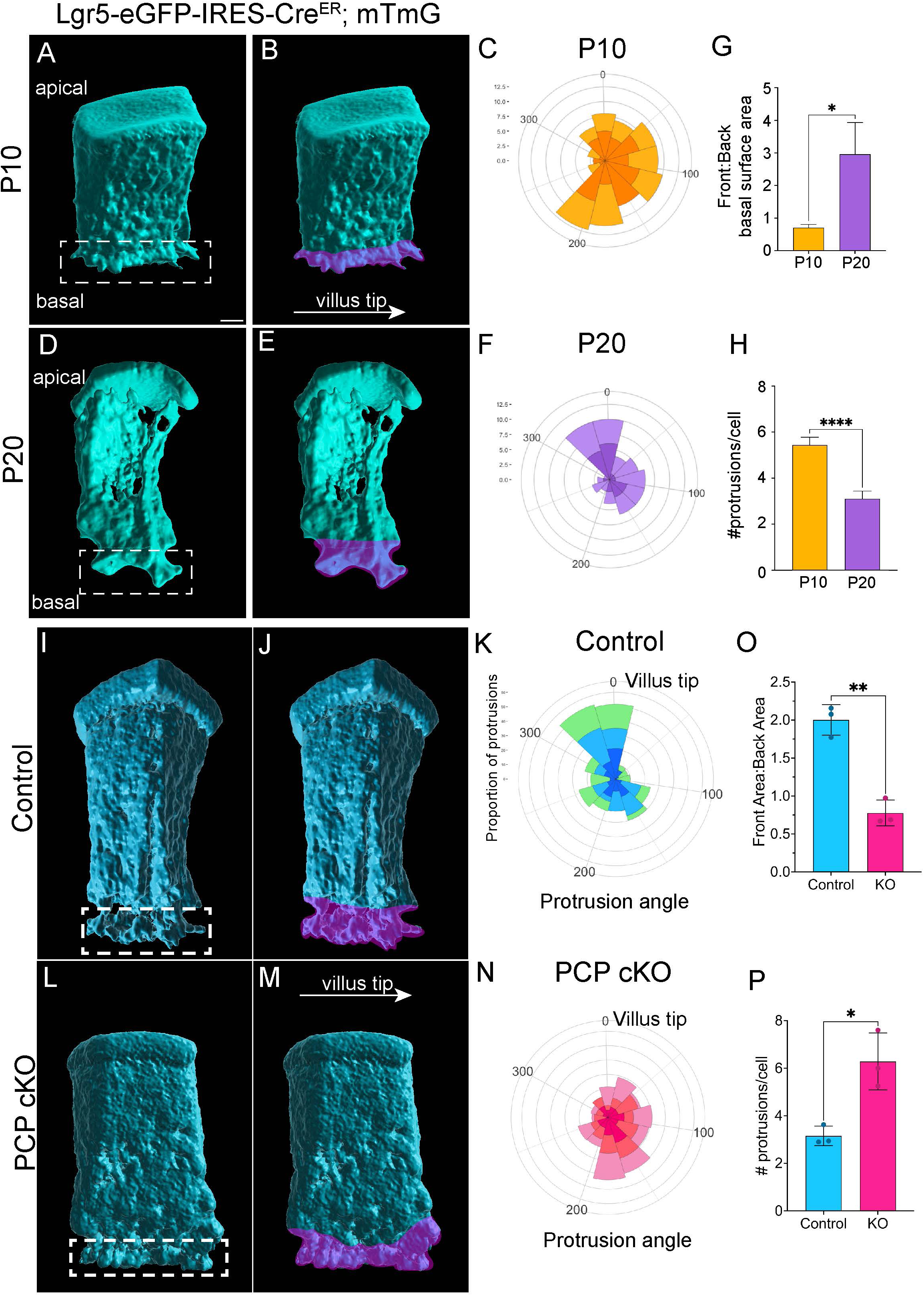
PCP orients basal protrusions to coordinate directional migration. A-D) Representative 3D rendered image of individual cells from Lgr5-EGFP-IRES-Cre^ER^; mTmG intestines at (A-B) P10. In B, a pseudocolored overlay marks the protrusive region. C) Rose plot illustrating the relative distribution of protrusion angles relative to villus tip (= 0°) at P10. D-E) Representative cell at P20. In E, a pseudocolored overlay marks protrusive region. F) Rose plot illustrating relative distribution of protrusion angles relative to villus tip at P20. G) Quantification of the front-to-back basal surface area between P10 and P20 enterocytes. H) Protrusion number per cell at P10 and P20. I-J) Representative cell in control P20. In J, pseudocolored overlay marks protrusive region. K) Rose plot illustrating relative distribution of protrusion angles relative to villus tip in control. L-M) Representative cell in PCP cKO (Vangl1/2 dKO) P20. In M, pseudocolored overlay marks protrusive region. N) Rose plot illustrating relative distribution of protrusion angles relative to villus tip in PCP cKO. O) Quantification of the front-to-back basal surface area between control and PCP cKO P20 enterocytes. P) Protrusion number per cell in control and PCP cKO P20 intestines. *, *p* < 0.05; **, *p* < 0.01; ****, *p* < 0.0001.

To test whether the migration defect in PCP cKO enterocytes is due to disrupted basal protrusion polarization, we utilized iMem-Mosaic^14^ in the Vangl1/2 dKO background to 3D reconstruct individual cKO cells and capture their protrusions at P20. Three-dimensional surface reconstructions of control and PCP cKO villar cells at P20 revealed striking morphological differences. In control cells, protrusions exhibited a clear front-to-back asymmetry, with basal protrusions oriented toward the villus tip (Fig 5I-K). In PCP cKO cells, basal protrusions were randomly oriented (Fig 5L-N). Quantitative analysis further revealed that front-to-back surface area ratio decreased significantly in PCP cKO cells (Fig 5O), and in fact, was reminiscent of P10 WT cells. Similarly, the number of protrusions was increased in PCP cKO cells compared to WT cells at P20 (Fig 5P). Thus, PCP signaling is required to orient protrusions toward the villus tip. Together, these data show that postnatal villus maturation involves the progressive polarization of basal protrusions, and that PCP signaling provides the cue that aligns these structures across cells. This cellular polarity forms the mechanical basis for the linear, non-overlapping villus ribbons characteristic of organized epithelial migration. When PCP is disrupted, protrusions do not orient, coherence breaks down, and ribbons maintain wavy and intersecting patterns.

### Polarized integrin localization is associated with linear migration patterns

Because basal protrusions engage the basement membrane, we next examined whether integrin localization changes during postnatal development, when migration becomes organized. In early postnatal intestines (P5, P10), ⍺6- and β4 integrin were distributed broadly across the basal surface of villus enterocytes, with little evidence of planar bias (Fig 6A-B,A’-B’). This localization became restricted over time (Fig 6A-D’), such that by P20, ⍺6- and β4 integrin were localized in discrete domains on the trailing edge of the basal side of each enterocyte (Fig 6D-D’). Line scan analysis across the basal surface of cells revealed a significant increase in ⍺6- and β4 integrin intensity in the trailing region of the cell between P5 and P20 (Fig 6E-F’). This is in contrast to phalloidin, which was consistently localized to the cell cortex at all time points examined (Fig 6G-G’). These results indicate that integrin planar polarizes progressively during villus maturation, coinciding with the emergence of directional protrusions and coherent epithelial migration.

**Figure 6.**
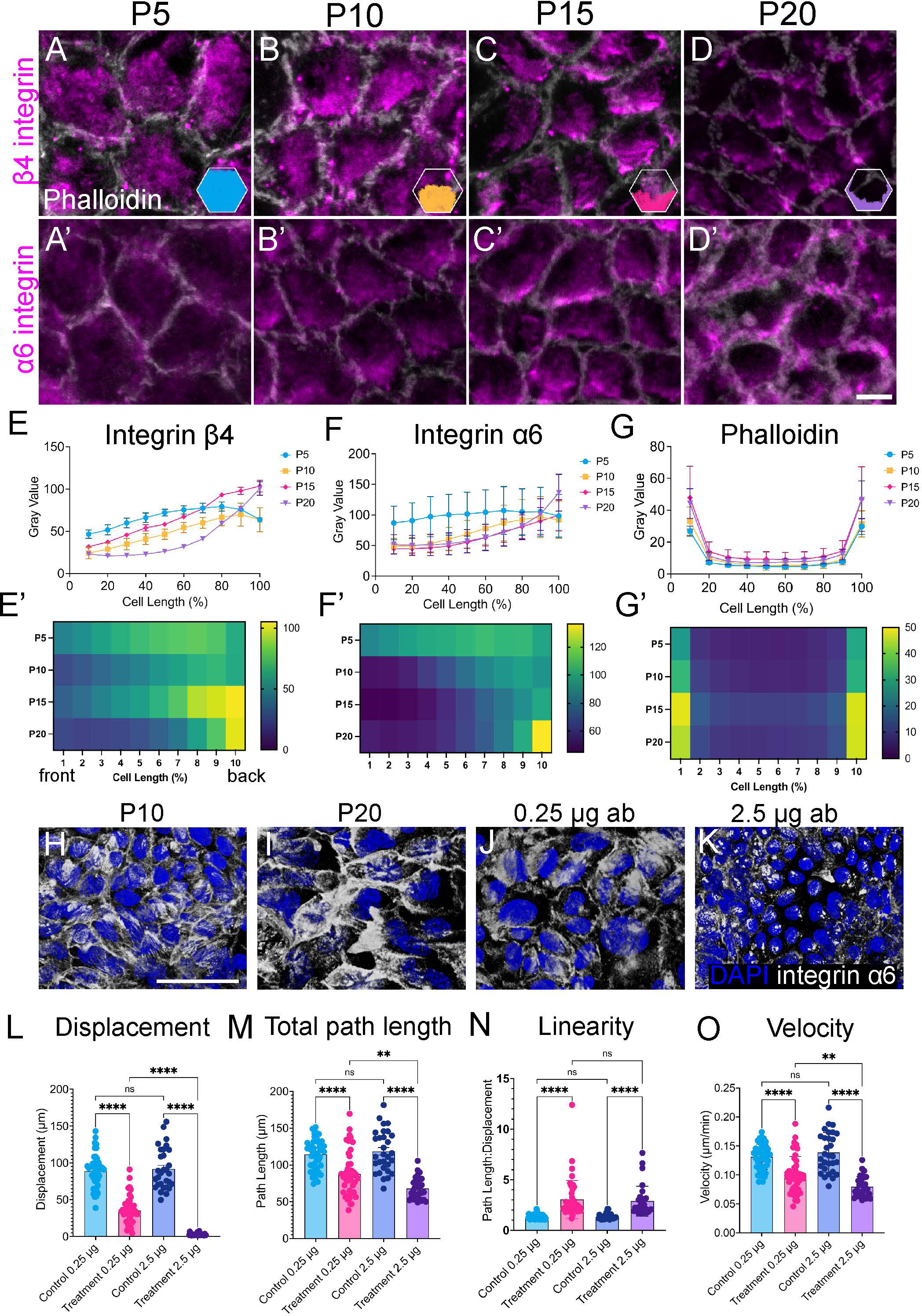
Integrin polarization reinforces coherent migration along the basement membrane. A-D’) Representative confocal images of β4 (magenta; A-D) and ⍺6 integrin (magenta; A’-D’) localization in villar enterocytes at P5, (A), P10 (B), P15 (C) and P20 (D). Phalloidin (white) marks cell boundaries. Scale bar, 5 µm. In A’-D’, an inset schematizes the localization pattern for each stage. E-G’) Line scan analysis of β4 integrin (E-E’), ⍺6 integrin (F-F’) and phalloidin (G-G’). H-I) ⍺6 integrin (white) localization in 2D enteroid monolayers derived from P10 (H) and P20 (I) intestines. J-K) Localization of ⍺6 integrin (white) in P20 2D enteroid monolayers treated with 0.25 µg (J) or 2.5 µg (K) ⍺6 integrin-blocking antibody. L-O) Quantification of cell migratory behavior in response to ⍺6 integrin-blocking antibodies.

To determine whether integrin engagement modulates migration behavior in our system, we manipulated adhesion strength in P20 enteroid monolayers. Interestingly, even in vitro, ⍺6 integrin localization was polarized to one edge to individual cells at P20 (Fig 6I). In contrast, cells within P10 monolayers did not polarize their ⍺6 integrin (Fig 6H). We reduced ⍺6-integrin binding using a blocking antibody. Low doses (0.25 µg) resulted in uniform localization of ⍺6- integrin along the basal surface of individual P20 cells (Fig 6J), while high doses (2.5 µg) collapsed ⍺6-integrin localization into plaques not associated with cell edges (Fig 6K). These data suggest that ⍺6-integrin activity can modulate its polarized localization patterns.

To test whether polarized integrin localization was necessary for directed migration of enterocytes, we tracked migration paths and velocity of cells in P20 monolayers treated with low or high doses of ⍺6-integrin blocking antibody at plating. While control cells collectively migrated in relatively linear paths (Fig 6L-N), the migration of cells treated with 0.25 µg ⍺6-integrin blocking antibody was less coordinated with neighbor cells, slower, and less processive (Fig 6L- O). In contrast, higher doses of ⍺6-integrin blocking antibody (2.5 µg) resulted in almost complete inhibition of cell migration (Fig 6L).

Together, these data identify integrin polarization as a developmental hallmark of villus maturation. As ⍺6/β4-integrins become spatially restricted along the basal surface, their engagement with the basement membrane fine-tunes the efficiency and coherence of epithelial migration, linking cell-intrinsic polarity to collective tissue dynamics.

### Loss of PCP weakens epithelial anchorage and distorts villus architecture

Because integrin polarity is refined during villus maturation, we next asked whether PCP signaling is required to maintain this organization and preserve epithelial attachment to the basement membrane. While the majority of the intestinal structure was intact in PCP cKO intestines, there were localized architectural defects characterized by branched structures that contained translocated crypts (Fig 7A-B, arrowheads; Fig 7J) and multiple villar branches (Fig 7B, J, dashed line). The translocated crypts were associated with gaps in the region above the muscle layers, where crypts normally reside (Fig 7B, asterisks). In each of these structures, there was one main villus branch that contained a mesenchymal core and extracellular matrix, and all of the epithelial branches were emanating from the main villus branch. In 2D cryosections, ⍺6 integrin appears as a continuous sheath underlying the epithelium in WT intestines (Fig 7C). Strikingly, within the ectopic branches of PCP cKO intestines, ⍺6 integrin localization was punctate instead of continuous, with a single collapsed punctum per cell (Fig 7D). Within crypts, entire structures were detached from the basement membrane, marked by the crypt-specific laminin ⍺2 (Fig 7E-F). Within ectopic epithelial branches in PCP KO villi, there was a complete lack of ECM (Fig 7G-H). Visualization of these structures in 3D whole mount tissue demonstrates the dramatic disorganization of the crypt-villus axis (Fig 7K-L).

**Figure 7.**
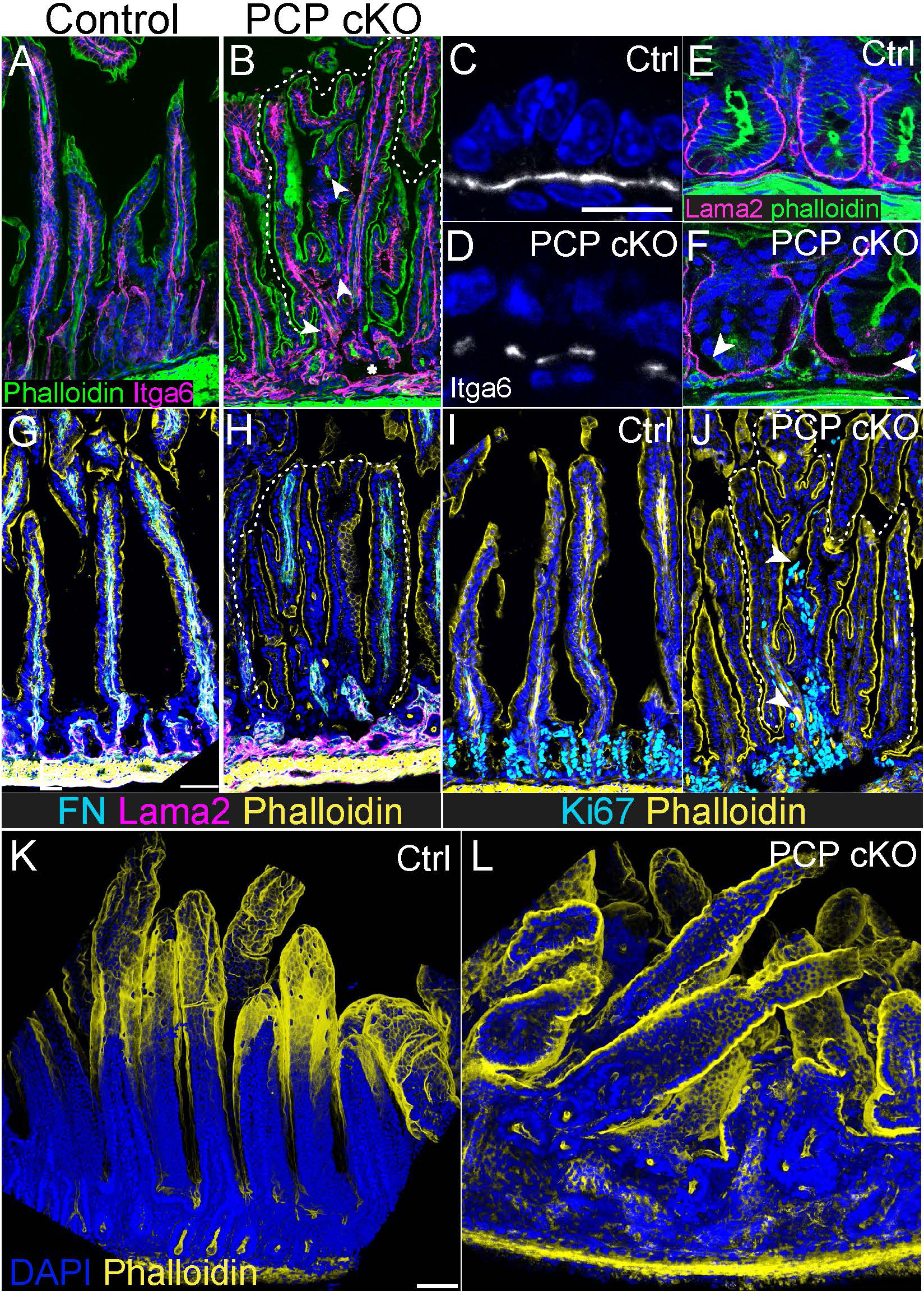
PCP maintains epithelial anchorage and preserves villus architecture. A-B) Cryosections from P20 control and PCP cKO intestines demonstrating architectural defects in PCP cKO, including branched villi (dashed line), crypt translocation (arrowheads) and gaps above the muscle layer (asterisk). Phalloidin is in green, and the basement membrane is marked by ⍺6 integrin (magenta). C-D) High magnification images of ⍺6 integrin staining. In PCP cKO villi, staining becomes punctate within branched villi. E-F) Immunostaining for crypt- specific basement membrane marker laminin ⍺2 (magenta) and phalloidin (green) highlights loosening of crypt epithelium from its underlying basement membrane (arrowheads). G-H) Fibronectin (cyan) and Lama2 (magenta) staining with phalloidin (yellow) demonstrates that villus branches (dashed line) do not contain detectable levels of EC. Translocated crypts are not associated with lama2 signal. I-J) Ki67 (cyan) and phalloidin (yellow) demonstrate the presence of translocated crypts (arrowhead) within branched villar regions (dashed line) of PCP cKO intestines. K-L) Whole mount staining of control (K) and PCP cKO (L) intestine for phalloidin (yellow) highlight the disorganization of the intestinal tissue in PCP cKO.

Together, these data show that villus deformation in PCP cKO tissue coincides with focal loss of integrin localization and basement membrane continuity. PCP signaling thus acts to stabilize epithelial anchorage, ensuring that the directional migration machinery established by protrusions and integrin polarization is transmitted uniformly across the villus surface. When PCP is compromised, these adhesive networks fail locally, producing the branching defects that disrupt villus architecture.

### Disrupted epithelial migration expands villus functional domains

Because epithelial migration along the villus establishes continuous ribbons of cells that differentiate in position-specific ways^15^, we asked whether the loss of migration coherence in PCP cKO intestines altered the spatial patterning of specialized functions (Fig 8A). We focused on the villus tip zone, where differentiation is terminal and positional identity is most sharply defined, reasoning that migration defects would have the strongest functional consequences at this final stage. In control tissue, the villus-tip specific enzyme adenosine deaminase (Ada) was sharply confined to the uppermost 10-15% of each villus (Fig 8B, D), whereas in PCP cKO villi, the Ada-positive domain expanded downward to encompass ∼30% of villus height (Fig 8C-D).

**Figure 8.**
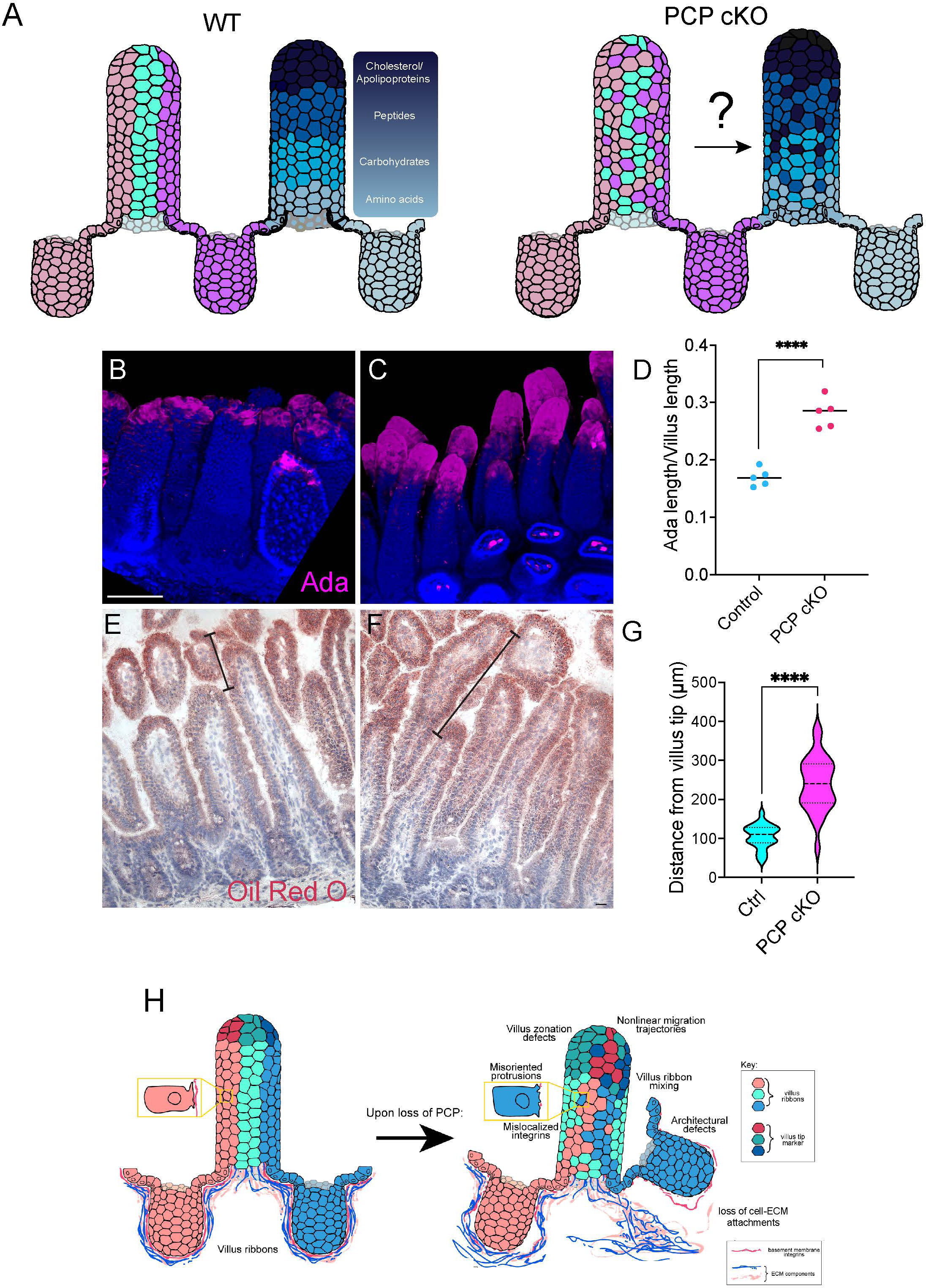
Disrupted migration coherence expands villus functional domains. A) Schematic illustrating the potential relationship between epithelial migration and villus zonation. In wild-type (WT), linear migration ribbons convey differentiating enterocytes upward along a stable basement membrane, maintaining discrete transcriptional and metabolic zones. B-C) Whole mount immunostaining of control (B) and PCP cKO (C) intestines stained for villus tip marker Ada (magenta). D) Quantification of Ada-positive area along the villus axis, expressed as fraction of villus height. E-F) Oil Red O staining following corn oil gavage, used to assess spatial distribution of lipid accumulation. High-intensity staining region is marked by bracket. G) Quantification of the Oil Red O enrichment zone, measured as the fraction of villus length occupied by high-intensity staining. H) Integrated schematic model. PCP signaling aligns basal protrusions and integrin polarity to generate coherent, linear migration ribbons that maintain positional identity along the villus. Loss of PCP decouples these axes, as misaligned protrusions and irregular basement membrane topology lead to nonlinear migration, disrupted villus architecture and disorganized zonation of gene expression and metabolic function.

Because this expansion reflected a loss of positional confinement rather than altered differentiation, we next examined whether functional zonation was similarly disrupted. Lipid absorption and chylomicron packaging are enriched in villus-tip enterocytes, forming one of the most spatially polarized metabolic programs in the intestine. We therefore administered a bolus of corn oil via gavage and sacrificed mice 20 minutes later. We then used Oil Red O staining as an independent readout of functional zonation, reasoning that the distribution of lipid droplets would reveal whether functional compartmentalization was preserved. In control villi, Oil Red O signal was enriched to the uppermost region of the villus, forming a narrow band of strong lipid accumulation at the villus tip (Fig 8E). In contrast, PCP cKO villi displayed a downward- expanded zone of enrichment that encompassed parts of the mid-villus (Fig 8F). Quantitative analysis revealed a > 2-fold expansion of the enriched lipid accumulation zone (Fig 8G). Thus, both molecular (Ada) and metabolic (Oil Red O) markers reveal a loss of villus tip confinement, indicating that PCP-driven linear migration is required to maintain the spatial fidelity of epithelial zonation.

### Integrated model: PCP aligns migration and zonation

Together, these findings reveal that PCP coordinates epithelial tissue polarity from the molecular to tissue scale. Under normal conditions, PCP orients basal protrusions and integrin attachments in register across neighboring cells, producing straight, coherent ribbons of migrating enterocytes that carry position-specific gene programs upward along the villus. This coordinated migration preserves functional coordination, ensuring that terminal differentiation and lipid-handling programs remain restricted to villus tip cells. When PCP signaling is lost, misaligned protrusions and disordered integrin polarity generate nonlinear trajectories that distort villus architecture and uncouple cell position from differentiation state. The schematic (Fig 8H) summarizes this hierarchy: PCP establishes a unified axis of epithelial motion linking molecular asymmetry to functional zonation. Loss of PCP decouples these layers, eroding the positional framework that sustains intestinal organization.

## Discussion

Tissue architecture and function are inextricably linked; the spatial organization of cells dictates how physiological processes are distributed and maintained. Here, we define a developmental mechanism that couples these levels through PCP. Using postnatal intestinal development as a model, we show that PCP signaling aligns the direction of epithelial migration, establishes mechanical continuity across cells, and thereby maintains functional zonation along the villus axis. Loss of PCP uncouples this hierarchy, disrupting protrusion orientation, integrin localization, and villus geometry, which in turn, expands villus tip gene and metabolic domains. These findings reveal that PCP signaling provides a physical framework through which local polarity cues are translated into tissue-scale patterning of function.

Based on previous studies in adult mice, villus ribbons appear to result from the coordination of oriented cell migration, cell boundary formation, and interactions with the extracellular matrix to form linear stripes along the villus^2,3,16^. However, it was not known how these characteristics are initially established and regulated. Because these patterning characteristics are not present until later postnatal stages, we hypothesized that villus cells migrate in broader, less polarized trajectories during early postnatal development and progressively refine their paths over time. Indeed, we find that actin-based basal protrusions become increasingly aligned toward the villus tip from P10 to 20, coincident with the maturation of linear ribbons and processive migration in enteroid monolayers. This acceleration in migration is consistent with previous studies that early postnatal self-renewal requires 11-14 days, while adult self-renewal requires 3-7 days^2,17^. These findings support a model in which the emergence of organized protrusive activity parallels the transition from stochastic to coherent migration. Understanding the factors that drive this transformation allows us to link the developmental establishment of migration patterns to their physiological significance.

Cell migration is essential for intestinal function, contributing to rapid epithelial turnover and the maintenance of the differentiation axis along the villus^11,15^. Despite the centrality of this process, the cues and mechanisms that provide directionality to epithelial movement have remained unknown. The PCP pathway is a conserved mechanism that governs directional migration across species, including convergent extension during gastrulation, neural crest cell migration, and axon guidance^18–22^. In the mammalian intestine, PCP has been implicated in embryonic elongation^23–25^, but its postnatal role was unknown. We identify a distinct postnatal function for PCP in aligning epithelial flow during villus maturation. Vangl2-GFP localization revealed crypt-enriched expression despite ubiquitous transgene activation, suggesting that the PCP complex may assemble preferentially in crypts to orient migration as cells exit the niche. Remarkably however, the resulting phenotype manifests in villi, where basal protrusions become misoriented in PCP mutants. While the link between compartment-specific effects requires further investigation, our findings are consistent with previous studies in the adult mouse cornea, where loss of PCP also leads to patterning defects due to alterations in epithelial migration trajectories^26^.

An important component of coordinated migration is the interaction between migrating cells and their substrate, the basement membrane and extracellular matrix (ECM), which provide traction and guidance. We find that basement membrane integrins become progressively planar polarized along the crypt-villus axis during the same period that protrusions align and migration becomes coherent. This developmental relocalization of ⍺6 and β4 integrins, together with actin polarization, suggests that integrins act as mechanical effectors translating PCP-dependent polarity into anisotropic traction. Blocking ⍺6 integrin binding *in vitro* disrupted both integrin localization and directional migration, demonstrating that integrin-mediated adhesion is necessary for processive movement. This echoes other systems such as *Drosophila* egg chamber rotation^27,28^, where collective migration remodels the ECM to generate feedback that reinforces directional movement. Once intestinal migration paths and ECM organization are established, it is unknown whether PCP functions to maintain their alignment and stability.

When PCP is lost, these polarity and adhesion systems collapse. ⍺6 integrin localization becomes discontinuous, laminin and fibronectin networks fragment, and epithelial attachments to the basement membrane weaken. The resulting loss of mechanical coherence leads to villus deformation and crypt translocation. These structural defects illustrate how PCP signaling integrates cell-intrinsic polarity with cell-ECM coupling to preserve epithelial architecture.

Importantly, these defects have functional consequences. In normal villi, cells migrate in straight, non-intersecting ribbons that maintain the spatial relationship between cell age, position, and differentiation state, allowing the formation of sharply confined transcriptional and metabolic zones. In PCP mutants, where migration becomes erratic and ribbons mix, this mapping collapses. Villus tip Ada expression extends down the villus axis, and Oil Red O staining reveals an expanded lipid uptake domain encompassing the mid-villus. Thus, coherent migration is necessary to preserve the positional fidelity of functional zonation. When the PCP-migration axis is disrupted, cells still differentiate, but their geographic organization no longer matches their transcriptional or metabolic identity.

Together, these findings define a hierarchical model linking PCP, integrin polarization, migration mechanics, and functional zonation. PCP aligns protrusions and adhesion complexes across neighboring cells to generate coherent epithelial motion. This directional migration maintains villus architecture and the spatial registry of specialized functions. Loss of PCP breaks this organizational structure; misoriented protrusions lead to disordered migration, weakened adhesion, architectural deformation and blurred zonation boundaries. PCP thus operates as a central organizer translating molecular asymmetry into tissue-scale order and functional compartmentalization.

Beyond intestinal biology, this mechanism exemplifies a general principle of epithelial design: tissue integrity depends on the alignment of polarity, adhesion and migration. The PCP-integrin axis we identify may operate in other renewing epithelia where coordinated migration sustains barrier function. Its disruption could underlie pathological remodeling and zonation loss in inflammatory or neoplastic disease. Altogether, our work reveals that PCP signaling not only builds epithelial architecture but reinforces it, ensuring that collective migration and spatial specialization remain mechanically and functionally in register.

## Materials and Methods

### Experimental Model and Details

#### Mice

All animal work was approved by the Institutional Animal Care and Use Committee (IACUC) of Yale University. Mouse strains used in this study were: VillinCreER^29^ (kindly gifted from Dr. Sylvie Robine, Institut Curie; JAX #020282), Lgr5-EGFP-IRES-CreER^30^ (Jackson Laboratories, #008875), Vangl2-GFP (kindly gifted from Dr. Toshihiko Fujimori, National Institute for Basic Biology (Shi et al., 2016)), Vangl1^fl/fl^ (Jackson Laboratories, #019518) ; Vangl2^fl/fl^ (Jackson Laboratories, #025174); Dvl2^fl/fl^ (Jackson Laboratories, #029061); iChr-Mosaic (Jackson Laboratories, #031302), iMb-Mosaic (Jackson Laboratories, #031301)^14^, ZO1-GFP^31^ (gift from Dr. Terry Lechler, Duke University), mTmG^32^ (Jackson Laboratories, #007676), and CD1 (Charles River). Mice were genotyped by PCR and both males and females were analyzed.

Sex-paired control and experimental littermates from both postnatal males and females 5 days or older were also used for analysis. All mice were housed and maintained in a barrier facility with a 12h-12h light-dark cycle (07:00-19:00 light) and ad libitum access to food.

### Monolayer Culture

Fresh monolayers were cultured as described previously with slight adjustments/modifications^33^. P10 and P20 mouse jejunum were dissected, cut into ∼2 cm pieces, and cut longitudinally open. The luminal surface was scraped with a glass coverslip to remove villi. The remaining tissue was incubated in 3 mM EDTA in PBS for 30 min at 4°C rotating. Tissue was lightly shaken in PBS to wash away the EDTA before being shaken in fresh PBS to release crypts. Individual crypts were selected for with a 70 µm cell strainer (Falcon #352350) and centrifuged at 300 x g for 5 min. The crypt pellet was washed once in DMEM (Thermo Scientific #11995065) before being resuspended in 2D attachment media (Recombinant Mouse EGF (Thermo Fisher Scientific #PMG8041), Recombinant Human Rspondin-1 (R&D Systems #4645-RS) or R-spondin conditioned media, LDN-193189 (Cayman #11802), CHIR99021 (Cayman #13122), Y- 27632 (Sigma #Y0503), Penicillin-Streptomycin-Glutamine (100X) (Thermo Fisher Scientific #10-378-016), Amphotericin B (American Bioanalytical #A2942)) and plated on 30 µl of 1:10 diluted Matrigel (Corning #356231) that had been solidifying at 37°C for at least 1 hour. After 2-4 hours, monolayers were washed twice with PBS before leaving them in conditioned media collected from L-WRN cells (kind gift from Thaddeus Stappenbeck, Washington University in St. Louis). The next day, the media was switched to ENR (EGF, Recombinant Human Noggin (R&D Systems #6057-NG), Rspondin-1). Media was changed every other day.

For a6 blocking experiments, 0.25 µg and 2.5 µg of a6 antibody (CD49f, BD Pharmingen #555734) or PBS was added to the media upon plating of crypts. Monolayers were allowed to grow for 3 days before live imaging overnight on the fourth day. After live imaging, monolayers were washed once with PBS and fixed in warmed 4% PFA in PBS for 10 min. They were then washed three times with PBS before being stored at 4°C.

### Organoid Culture

Organoids were generated as stock for later use in making monolayers. Organoids were cultured as described previously^34^ using conditioned media collected from L-WRN cells. Fresh organoids were generated by isolating individual crypts as previously described for monolayer culture. Crypts were centrifuged at 300 x g for 5 min, before being resuspended in Matrigel and plated in 40 µL domes. The Matrigel domes were solidified at 37°C for 15-20 min before L-WRN media was added. Media was changed every other day.

Existing organoid stock was passaged every 5-7 days. Organoids were incubated in TrypLE Express Enzyme (Falcon #12604013) for 2 min at 37°C before mechanically disrupting them via pipetting up and down. TrypLE was inactivated through dilution with an equal amount of DMEM. The fragmented organoids were centrifuged at 300 x g for 3 min, washed again with DMEM, spun down again at 500 x g for 3 min, and either plated in Matrigel domes to continue the organoid stock or resuspended in L-WRN media supplemented with EGF and Y-27632 and plated on 1:10 Matrigel to make monolayers. After 2-4 hours, media was changed to ENR. Media was changed every other day.

### CreER induction

#### Lgr5-EGFP-IRES-CreER; mTmG

Lgr5-EGFP-IRES-CreER mice were crossed to mTmG -/- mice to generate Lgr5-EGFP-IRES-CreER; mTmG +/- mice. To visualize villus ribbons and basal protrusions at various time points during postnatal development, mosaic expression of mGFP was induced in intestinal stem cells by administering 50 µg tamoxifen (Sigma-Aldrich #T5648) in corn oil (Sigma-Aldrich #C8267) via intragastric injection at postnatal day 1 or 2 (P1 or P2). Tissue was collected at P10, P12, P14, P16, P18, and P20.

#### PCP cKO (VillinCre^ER^; Dvl2 fl/fl; iChr-Mosaic, VillinCre^ER^; Vangl1^fl/fl^; Vangl2^fl/fl^; iMb-Mosaic, VillinCre^ER^; Dvl2 fl/fl, VillinCre^ER^; Vangl1^fl/fl^; Vangl2^fl/fl^)

VillinCre^ER^; Dvl2 fl/fl; *iChr-Mosaic* mice were used to visualize villus ribbons upon conditionally deleting PCP genes. VillinCre^ER^; Vangl1^fl/fl^; Vangl2^fl/fl^; *iMb-Mosaic* mice were used to visualize protrusion dynamics upon conditionally deleting PCP genes. VillinCre^ER^; Dvl2 fl/fl; *iChr-Mosaic* mice were used to visualize villus ribbon dynamics upon conditional deletion of PCP genes. To induce deletion of PCP genes (Dvl2, Vangl1, and Vangl2), 200 µg tamoxifen in corn oil was administered via intragastric injection at P2. Same-sex control littermates were injected with the same tamoxifen injection scheme. Tissue was collected at P20.

### Tissue Preparation

Mouse jejunum from postnatal day 20 (P20) mice was isolated and embedded in Tissue-Tek O.C.T. Compound (OCT; Sakura Finetek USA #4583) and frozen on dry ice. Frozen OCT blocks were sectioned at 8 µm thickness using a cryostat.

For experiments requiring paraformaldehyde fixation, postnatal mouse jejunum was first cut into ∼2 cm pieces and flushed with 1xPBS. Tissue was then flushed with 5-10 mL of 4% paraformaldehyde (PFA; Sigma-Aldrich #158127) in PBS with 1mM CaCl2 before being left in 4% PFA in PBS with 1 mM CaCl2 overnight. The next day, tissue was washed with PBS three times. For prefixed tissue cryosections, PFA fixed P20 tissue was submerged in 20% sucrose in PBS overnight at 4°C for sucrose protection. Sucrose-protected tissue was then embedded and frozen on dry ice before being stored in -80°C. Frozen OCT blocks were sectioned at 8 µm thickness using a cryostat. For vibratome samples, PFA fixed P10, P12, P14, P15, P16, P18, and P20 mouse jejunum were stored in PBS containing 0.2% PFA at 4°C. To create 150 µm vibratome sections, fixed tissues were first embedded in 2.5% agarose (low-gelling temperature, Sigma-Aldrich #A9414). Tissue was cut at speed 3.5 and oscillation 4 with the Precisionary Compresstome (Model: VF-310-0Z). Intestinal rings were stored in PBS with 0.2% PFA at 4°C for up to two weeks prior to use.

For whole mount tissues, P20 mouse jejunum was first dissected into ∼2 cm pieces and then cut longitudinally to expose the epithelium. Flat intestinal pieces were pinned down on PDMS plates and fixed in 4% PFA in PBS with 1mM CaCl2 at room temperature (RT) for two hours. After fixation, whole mount tissues were washed with PBS three times and stored at 4°C in PBS containing 0.2% PFA.

### Immunofluorescence

For cryosection staining, sections were fixed for 8 min using 4% PFA in PBS and then washed with PBS-T containing 0.2% Triton X-100 (AmericanBIO #AB02025). Sections were incubated in blocking buffer containing 3% BSA (Sigma-Aldrich #A9647) and 5% NDS (Jackson ImmunoResearch Labs #017-000-121) in PBS-T for at least 15 min. Then, primary antibodies diluted in blocking buffer were added to sections for 15 min-1 hour at RT or overnight at 4°C. Sections were then washed in PBS-T for 5 minutes three times. After PBS-T washing, secondary antibodies diluted 1:200 in blocking buffer were applied onto sections for 10 min.

Sections were washed again in PBS-T for 5 minutes three times. After PBS-T washing, sections were mounted in antifade medium, consisting of 90% glycerol (JT Baker #2136-01) in PBS plus 2.5 mg/ml *p*-Phenylenediamine (Acros Organics #417481000).

For whole mount and vibratome staining, PFA fixed tissues were incubated in blocking buffer overnight or for 2 hours at RT on a nutating mixer, respectively, followed by primary antibody incubation for 2-3 days at 37°C. Tissues were washed three times in PBS-T containing 0.2% Triton X-100 then incubated in secondary antibodies for 1 day at 37°C. Then, PBS-T washes were repeated prior to tissue clearing with Ce3D medium overnight at RT. Tissues were imaged in Ce3D clearing medium^35–37^ or FocusClear (Cedarlane #FC-101).

For monolayer staining, fixed monolayers were incubated in PBS-T for 10 min before being left in blocking buffer for 30 min at RT. Primary antibodies diluted in blocking buffer were added for 2 hours at RT before being washed three times with PBS-T. Secondary antibodies were then added for 30 min. Monolayers were then washed three times in PBS-T with a final wash in PBS. Monolayers were imaged in PBS.

The following primary antibodies were used: rabbit anti Ki67 (Abcam #ab15580), rat anti Epcam (Biolegend #118202), rat anti CD44v6 (Thermo Fisher #BMS145), rabbit anti ZO1 (Invitrogen #61-7300), chicken anti GFP (abcam #ab13970), rat anti integrin a6 (BD Pharmingen #567005), rat anti integrin b4 (BD Pharmingen #553745), rabbit anti fibronectin (END Millipore #AB2033), and rat anti laminin a2 (Santa Cruz #sc-59854). Tissue sections were imaged on an upright Zeiss AxioImager with Apotome 2 attachment and Zeiss AxioCam 506 mono camera using Zen software (v3.0; Zeiss). Objectives used were Plan Apochromat 10x/0.45 air, 20x/0.8 air, 40x/1.3 oil, and 63x/1.4 oil. Whole mount tissues, vibratome sections, and 2D enteroid monolayers were imaged on an inverted Leica Stellaris 5 confocal laser scanning microscope using Diode 405, a white light laser and LAS-X software (v4.6.1; Leica). Objectives used were HC PL APO CS2 10x/0.40 dry, 40x/1.10 water, 40x/1.30 oil, 63x/1.40 oil and HC FLUOTAR L VISIR 25x/0.95 water.

### Time-lapse image acquisition and microscopy

Imaging of 2D enteroid monolayers was performed on an inverted scanning confocal microscope. Monolayers were imaged in a humidity and temperature-controlled chamber at 37°C and 5% CO2. A 40x/1.3 oil objective (Leica) was used and the pinhole was opened to 2 to prevent further photodamage.

### RNA sequencing analysis

RNA-sequencing analysis was performed on a publicly available dataset^7^ (GSE109054). DESeq2 analysis was performed to compare expression of candidate genes between P6 and P10 samples.

### Quantification and Statistical Analysis

#### Image analysis

Quantifications of villus ribbons and protrusions were performed in Imaris (v10.2.0; Bitplane) on volume rendered images of intestinal vibratomes and whole mounts. For villus ribbons, nuclei were detected using the Imaris Spots module. The x, y coordinates were extracted and used to determine the proportion of cells belonging to Ribbon A vs. Ribbon B. For protrusions, Imaris Surfaces was used to 3D reconstruct individual cells to mask out the cell of interest. The cell was reoriented using Imaris’ Free Rotate for better positioning when it was reopened in Fiji/ImageJ (v 2.16.0/1.54p). After identifying the neck region of the cell (relatively apical) and the basal protrusions, protrusions were identified with the use of a previously published macro^11^. Protrusion angle, length, and number were identified manually. This data was inputted into R (version 4.5.1 (2025-06-13)) to generate rose plots.

For quantification of villus ribbon boundary linearity, Fiji/ImageJ was used to measure the perimeter length of cell membranes at the border of villus ribbons. This measurement was compared to the corresponding shortest path length between the bottom and top of the villus ribbon as a proxy to determine the linearity of cell migration within that ribbon. Only ribbons derived from clonal crypts were quantified.

Monolayer tracking of nuclei was done by first using Cellpose in Aivia (v14.1.0; Leica) to segment and mask out the nuclei in max projections of the monolayer time-lapses. This mask was then used to perform Cell Tracking in Aivia.

Statistical analysis was performed using GraphPad Prism

## Acknowledgments

We thank all members of the Sumigray lab for critical discussions and Maria Figetakis for careful mouse colony management. We thank Dr. Toshihiko Fujimori for the Vangl2-GFP mouse line, Dr. Danelle Devenport, Dr. Terry Lechler and Dr. Valentina Greco for reagents and critical discussions and feedback. We thank Dr. Caroline Hoppe for help with 3D reconstructions and statistical analysis. We would also like to acknowledge the Center for Cellular and Molecular

Imaging (CCMI) at Yale and Daniel Miranda (AIVIA, Leica) for their resources and expertise in image analysis. This work was funded in part by NIH R35GM150645 to KS, a Chen Innovation award from the Yale Stem Cell Center to KS, and a Yale Stem Cell Center Lo Fellowship to RFL.

